# FastID: Extremely Fast Forensic DNA Comparisons

**DOI:** 10.1101/173666

**Authors:** Darrell O. Ricke

**Affiliations:** Bioengineering Systems & Technologies, Massachusetts Institute of Technology Lincoln Laboratory, Lexington, MA USA

**Keywords:** DNA forensic, identification, mixture analysis, familial search

## Abstract

Rapid analysis of DNA forensic samples can have a critical impact on time sensitive investigations. Analysis of forensic DNA samples by massively parallel sequencing is creating the next gold standard for DNA forensic analysis. This technology enables the expansion of forensic profiles from the current 20 short tandem repeat (STR) loci to tens of thousands of single nucleotide polymorphism (SNP) loci. A forensic search scales by the product of the number of loci and the number of profile comparisons. This paper introduces a method (FastID) to address the need for rapid scalable analysis of DNA forensic samples (patent pending)[1]. FastID can search a profile of 2,500 SNP loci against 20 million profiles in 5.08 seconds using a single computational thread on a laptop (Intel i7 4.0 GHz).

## I. INTRODUCTION

Advances in High Throughput Sequencing (HTS) enables Massively Parallel Sequencing (MPS) of DNA forensic samples[2]. MPS enables sequencing of Short Tandem Repeats (STRs) and/or Single Nucledotide Polymorphisms (SNPs)[3]. Law enforcement and forensics communities in the United States currently use a forensic panel called the Combined DNA Index System (CODIS). This forensics panel is composed of 20 STR loci. MPS panels can expand CODIS by tens of thousands of SNP loci. This increase in panel size will enable significantly expanded analysis potential with increased confidence. The FBI NDIS database currently has over 16 million STR profiles[4]. Searching a STR profile of 20 loci times 2 alleles for each loci times 16 million profiles requires 20 X 2 X 16 million comparisons – complexity order of O(N x M). Expanding the size of panels to 20,000 loci represent an increase of the number of required comparisons by a factor of 1 million over the current CODIS panel. For DNA forensics, this is a relevant computational issue to address.

Forensic capabilities, not currently achievable by the current CODIS STR panel, are created by adding SNPs to DNA forensic panels. Panels of SNPs have been created for enhanced DNA mixture analysis[5], enhanced kinship identification[6], biogeographic ancestry (BGA) predictions[7], phenotype/externally visible traits (EVTs) predictions[8], and the ability to analyze trace DNA samples. Also, additional Y chromosome STRs have been used for surname predictions[9]. To determine if a person is present in a mixture, the probability of random man not excluded, P(RMNE), is calculated[5]. It is broadly-expected that expansion of DNA forensic panels will be widely adopted in the future.

The FastID method was created to enable rapid searching of forensic panels with large numbers of loci. FastID uses a new data encoding that enables parallel comparisons of SNPs using standard computer hardware instructions of exclusive OR (XOR), logical AND, and population count (pop count). FastID can search a profile of 2,500 SNPs against a database of 20 million profiles in 5.08 seconds on a laptop (Intel i7) 4.0GHz using a single computational thread.

As with current STR-based search techniques, FastID can search a DNA profile from a suspect (e.g., a ‘reference’ profile) and match it against all available crime scene DNA samples (‘forensic’ profiles) and known DNA reference profiles. It can also take as input a crime scence DNA sample (forensic profile) and match it against all known DNA reference profiles and DNA samples from different crime scenes. In other words, FastID implements “identity searching” (reference profiles versus reference profiles) and “mixture analysis” (forensic profiles versus reference profiles or reference profiles versus forensic profiles).

## II. METHODS

### A. SNP Alleles Encoding

The majority of SNPs have two alleles (alternative forms of a gene). In a population, the more frequent allele is referred to as the major allele (M) and the less frequent allele as the minor allele (m). SNPs with low minor allele frequencies across a wide variety of ethnicities are optimal for DNA mixture analysis[5, 10]. Let p represent the major allele frequency and q the minor allele frequency (p + q = 1). For N loci, a typical individual should have Np^2^ major alleles (MM), 2Npq loci with one major and one minor allele (mM), and Nq^2^ loci with two minor alleles (mm). FastID encodes both SNP alleles as a single binary bit for rapid comparisons (Table 1). Vectors of SNPs can be mapped onto hardware representations as bit arrays (or integers). The sign bit of integers is not used to avoid automatic conversions of encoded bits to the two’s complement encodings of integers, which impacts the encoded SNPs.

**Table 1.** Binary Encoding of SNP Alleles

| Alleles | Binary Encoding |
| --- | --- |
| MM | 0 |
| Mm | 1 |
| mm | 1 |

### B. Reference Comparisons

Two reference samples can be compared with the logical exclusive OR (XOR) operation. Any differences between the profiles will result in one bits set for each SNP with a difference between the two samples. The total number of SNPs with differences can be tallied with the hardware population count (pop. count) instruction. Only 2 hardware instructions are needed for each block of SNPs to be compared. On a 64-bit computer each block consits of 60 SNPs to avoid using the integer sign bit. Ordered SNPs are encoded as hexidecimal strings for reference and mixture profiles as input to FastID. A final population count sum of zero represents a perfect match between two reference samples. A final sum greater than zero represents the number of SNPs with different minor allele encodings. Table 2 illustrates two reference profiles for the same individual with each profile illustrating one “dropped” (e.g., missing) minor allele (red) or, alternatively, “dropped in” alleles. The probability of random man not excluded, P(RMNE), calculation provides insight into the signficance of the comparison[5]. When genotypes for individuals can be predicted, the forensics community prefers the Bayesian forensic Likelihood Ratio (LR) calculation over P(RMNE) based on allele frequenceis. The P(RMNE) threshold minimum of 1e-6 is used to obtain search warrents and 1e-9 for criminal prosecution. Lower P(RMNE) can be obtained with larger SNP panels.

**Table 2.** Comparison of Individual 1 Replicate Samples (-a & -b) – 1 in 6.71 million chance of random match (1/1.491e-7)

| Sample | Minor Alleles encoded as Hexadecimal digits |
| --- | --- |
| Individual-1-a | 02000000004000000004040010000002000000000800000200000000 |
| Individual-1-b | 02000000000000000004040010000002000000000800000200000040 |
| XOR | 000000000040000000000000000000000000000000000000000000040 |
| Population count | 2 |
| P(RMNE) | 1.491e-07 |

### C. Mixture Comparisons

A forensic profile may contain a mixture of DNA from more than one contributor. In FastID, mixture analysis is implemented by taking the result of an exclusive OR (XOR) of a reference profile and a forensic profile combined with a logical AND to the reference profile followed by the population count instruction. The logical AND operation identifies only differences on the encoded minor alleles in the reference profile. This three-instruction combination restricts the population count sum to reflect the number of minor alleles in the reference profile that are not found in the forensic profile. The final sum of the population count represents the number of SNPs with minor alleles in the reference profile that are not present in the forensic profile. Table 3 illustrates the comparison of a reference profile with a forensic profile in which only one SNP with a minor allele is discordant with the forensic profile. This represents a very close match between the reference profile and the forensic profile with a P(RMNE) of 2.74e-7. Table 4 illustrates the comparison of a different reference profile with the same forensic profile; Individual-2 has 14 SNPs with minor alleles that are discordant with the forensic profile and a P(RMNE) value of only 0.0785. This represents a very poor match between the second reference profile and the forensic profile for this small set of SNP loci comparisons. Roughly 1 in 12.7 (1/0.0785) random individuals will likely match this forensic profile by random chance alone.

**Table 3.** Comparison of Individual 1 (-a) with Mixture 3 – 1 in 3.65 million that this match is random

| Sample | Minor Alleles encoded as Hexadecimal digits |
| --- | --- |
| Individual-1-a | 02000000004000000004040010000002000000000800000200000000 |
| Mixture-3 | 32288040038100200804462a5000568220205c485c38048398000440 |
| XOR | 3028804003c100200800422a4000568020205c485438048198000440 |
| AND | 000000000040000000000000000000000000000000000000000000000 |
| Population count | 1 |
| P(RMNE) | 2.740e-07 |

**Table 4.** Comparison of Individual 2 with Mixture 3 – 1 in 12.7 that this is a random individual (e.g., not in mixture)

| Sample | Minor Alleles encoded as Hexadecimal digits |
| --- | --- |
| Individual-2 | 06001440004808200000202000000006010000086000420000000003 |
| Mixture-3 | 32288040038100200804462a5000568220205c485c38048398000440 |
| XOR | 3428940003c908000804660a5000568421205c403c38468398000443 |
| AND | 04001400004808000000200000000004010000002000420000000003 |
| Population count | 14 |
| P(RMNE) | 0.0785 |

*This material is based upon work supported under Air Force Contract No. FA8721-05-C-0002 and/or FA8702-15-D-0001. Any opinions, findings, conclusions or recommendations expressed in this material are those of the author(s) and do not necessarily reflect the views of the U.S. Air Force.*

### D. FastID

FastID has been implemented as a single threaded C program. FastID takes two files as input: query profiles and target profiles, both with hexadecimal SNP encodings.

### E. In silico datasets

An *in silico* dataset was used to create target databases based on population SNP minor allele frequencies[11].

### F. Benchmark system

The SAGER laptop has two Intel 4.0 GHz i7 6700K CPUs with 4 cores each, 64 GB RAM, and RAID 1 512GB Samsung 850 EVO Series SATA3 Solid State Disks (up to 540 MB/sec sequential read).

## III. RESULTS

The total runtime for FastID includes both the time to read in the query and target profiles plus the FastID algorithm comparison times. The time to read the profiles scales with the number of SNPs and profiles being compared. Figure 1 illustrates the single Intel 4.0 GHz i7 6700K CPU thread runtime performance for FastID. Each plot represents different forensic SNP panel sizes of 500, 1,000, 2,500, and 5,000 SNPs. Each line represents different numbers of query profiles that are compared to target databases of 1 million, 5 million, 10 million, 15 million, and 20 million profiles. The FastID algorithm search times for one query profile can be estimated by subtracting the runtimes of a single query from the runtimes of 100 queries and dividing the result by (100-1=99) as illustrated in Figure 2. Figure 2 also illustrates the total number of SNP comparisons for the search times shown. The complexity of FastID comparisons for one query is N SNP comparisons times M target profiles or O(N x M).

**Figure 1.**
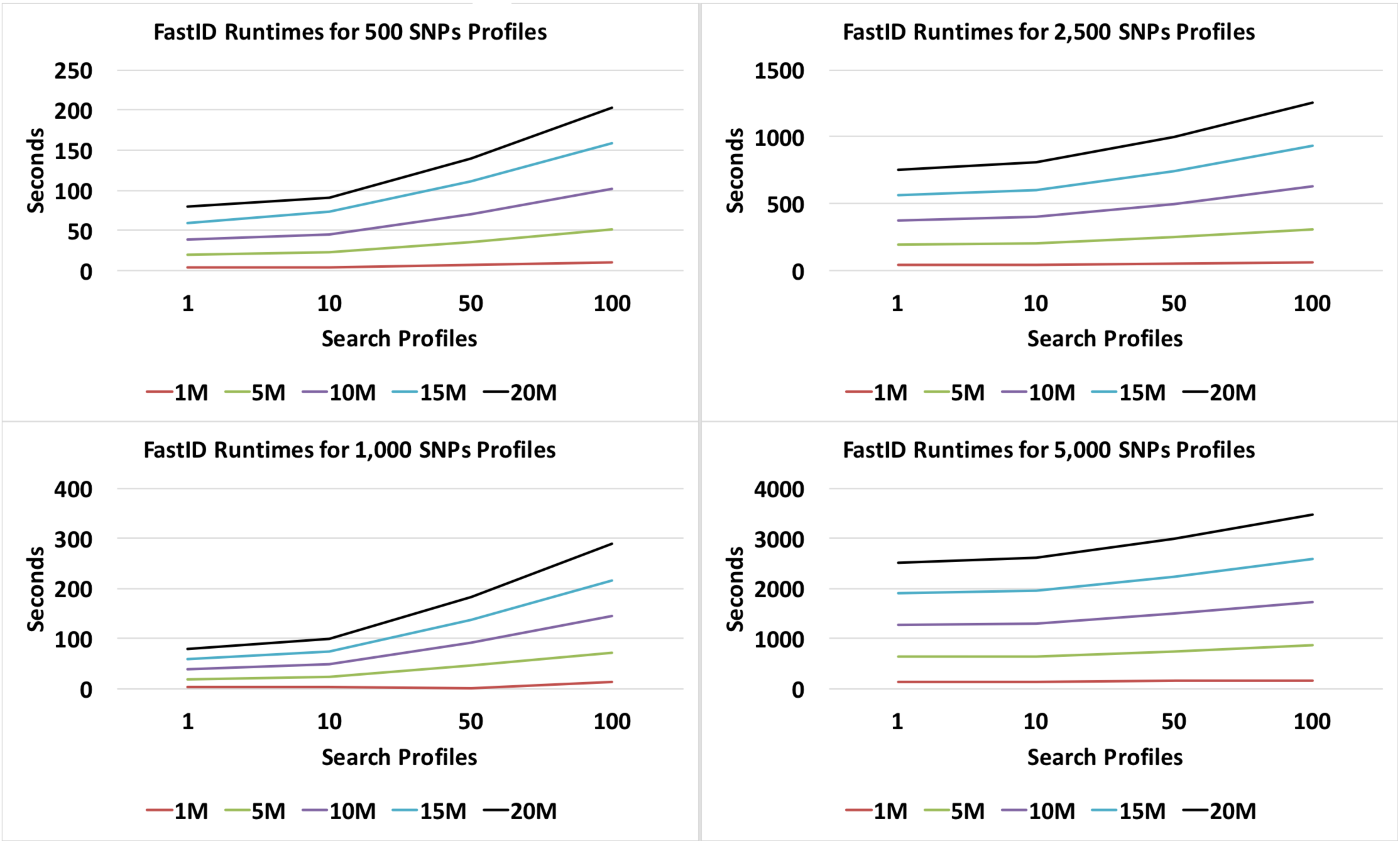
FastID Runtimes on 2.3GHz Xeon 2698 v3 CPU for 500, 1000, and 2500 SNPs for 1, 10, 50, and 100 queries against reference databases of 1M, 5M, 10M, and 20M profiles.

**Figure 2.**
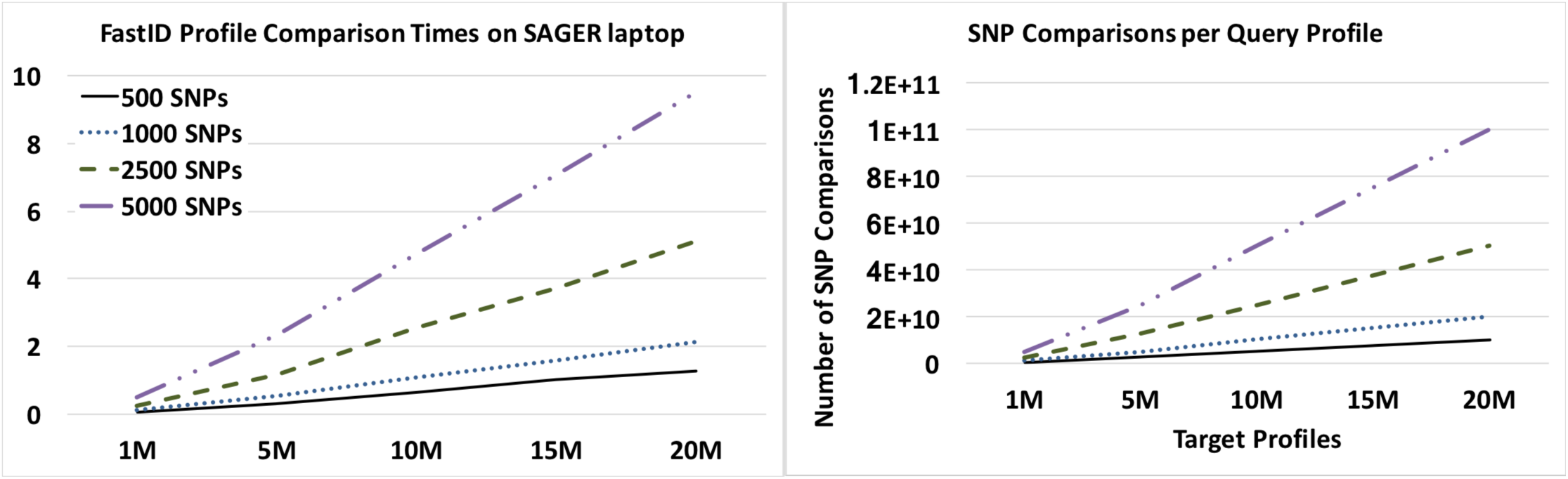
FastID Profile Comparison Times on 4.0 GHz i7 6700K CPU with single thread and number of SNP comparisons

FastID identifies perfect matches for query profiles if they are contained within the set of targets with a resulting value of zero. Nonzero results indicate the number of query minor allele mismatches to target profiles. The significance of match results can be evaluated by P(RMNE) value.

## IV. DISCUSSION

A profile search with 20 (formerly 13) STR loci against reference databases is the current forensic standard. The search time increases with the number of loci being compared times the number of profile comparisons O(N x M). Currently, processing of large numbers of STR profile comparisons requires larger computing systems and even entire data centers for organizations such as the FBI. Expanding the current STR panels to much larger SNP panels creates a potential computational bottleneck for processing larger numbers of samples against a database of 10 to 20 million profiles. FastID was developed as a solution to the computational bottleneck created when forensic MPS panels are increased from 20 to 2,000 or even 20,000 loci. Example FastID search times are illustrated in Figures 1 and 2. FastID detects identity matches (identification). FastID enables mixture analysis of complex mixtures with rapid comparisons of references to mixture profiles and forensic samples to references (Table 4).

## V. DISTRIBUTION STATEMENT

*A.* Approved for public release: distribution unlimited.

## REFERENCES

[1]. D. O. Ricke. Inventor; Massachusetts Institute of Technology, assignee. “DNA Mixtures from One or

[2]. D. O. Ricke, M. Petrovick, J. Bobrow, T. Boettcher, C. Zook, J. Harper, et al., “Human CODIS STR Loci Profiling from HTS Data,” Technologies for Homeland Security (HST), 2016 IEEE International Symposium on, 2016.

[3]. (2016). Illumina ForenSeq. Available:http://www.illumina.com/areas-of-interest/forensic-genomics/forensic-analysis-methods/snp-stranalysis.html

[4]. (2017). FBI CODIS -NDIS Statistics. Available: https://www.fbi.gov/services/laboratory/biometricanalysis/codis/ndis-statistics

[5]. J. Isaacson, E. Schwoebel, A. Shcherbina, D. Ricke, J. Harper, M. Petrovick, et al., “Robust detection of individual forensic profiles in DNA mixtures,” Forensic Science International: Genetics, vol. 14, pp. 31–37, 1//2015.

[6]. A. Shcherbina, D. O. Ricke, E. Schwoebel, T. Boettcher, C. Zook, J. Bobrow, et al., “KinLinks: Software Toolkit for Kinship Analysis and Pedigree Generation from HTS Datasets,” Technologies for Homeland Security (HST), 2015 IEEE International Symposium on, 2016.

[7]. R. Kosoy, R. Nassir, C. Tian, P. A. White, L. M. Butler, G. Silva, et al., “Ancestry Informative Marker Sets for Determining Continental Origin and Admixture Proportions in Common Populations in America,” Human mutation, vol. 30, pp. 69–78, 2009.

[8]. W. Branicki, F. Liu, K. van Duijn, J. Draus-Barini, E. Pośpiech, S. Walsh, et al., “Model-based prediction of human hair color using DNA variants,” Human Genetics, vol. 129, pp. 443–454, 2011.

[9]. M. Gymrek, A. L. McGuire, D. Golan, E. Halperin, and Y. Erlich, “Identifying Personal Genomes by Surname Inference,” Science, vol. 339, p. 321, 2013.

[10]. L. Voskoboinik and A. Darvasi, “Forensic identification of an individual in complex DNA mixtures,” Forensic Science International: Genetics, vol. 5, pp. 428–435.

[11]. B. S. Helfer and D. O. Ricke, “SNP Identity Searching Across Ethnicities, Kinship, and Admixture,” unpublished.

